# AFF-1 fusogen can rejuvenate the regenerative potential of adult dendritic trees via self-fusion

**DOI:** 10.1101/062679

**Authors:** Veronika Kravtsov, Meital Oren-Suissa, Benjamin Podbilewicz

**Affiliations:** Department of Biology, Technion-Israel Institute of Technology, Haifa 32000, Israel

**Author notes:** These authors contributed equally to this work (Veronika Kravtsov, Meital Oren-Suissa).

**Keywords:** AFF-1, dendritic remodeling, *Caenorhabditis elegans*, regeneration, selffusion

## Abstract

The aging brain undergoes structural changes, affecting brain homeostasis, neuronal function and consequently cognition. The complex architecture of dendritic arbors poses a challenge to understanding age-dependent morphological alterations, behavioral plasticity and remodeling following brain injury. Here, we use the PVD polymodal neurons of *C. elegans* as a model to study how aging affects neuronal plasticity. Using confocal live imaging of *C. elegans* PVD neurons, we demonstrate age-related progressive morphological alterations of intricate dendritic arbors. We show that insulin/IGF-1 receptor mutations (*daf-2*) fail to inhibit the progressive morphological aging of dendrites and do not prevent the minor decline in response to harsh touch during aging. We uncovered that PVD aging is characterized by a major decline in regenerative potential of dendrites following experimental laser dendrotomy. Furthermore, the remodeling of transected dendritic trees via AFF-1-mediated self-fusion can be restored in old animals by DAF-2 insulin/IGF-1 receptor mutations, and can be differentially reestablished by ectopic expression of AFF-1 fusion protein (fusogen). Thus, AFF-1 fusogen ectopically expressed in the PVD and mutations in DAF-2/IGF-1R, differentially rejuvenate some aspects of dendritic regeneration following injury.

**Summary statement:** Ectopic expression of AFF-1 fusogen or low activity of IGF-1R/DAF-2 rejuvenate the regeneration potential of dendrites following injury in aging C. *elegans*

## Introduction

Aging is the primary risk for neuronal diseases and general cognitive decline in humans (Yankner et al., 2008), yet, our understanding of the process of neuronal aging at the molecular and cell biological levels is still limited. In particular, very few studies have investigated the fate of complex dendritic arbors during aging, and their regenerative capacity following brain injury or trauma. Signaling via the DAF-2 Insulin/IGF-1 receptor is the most prominent and conserved pathway that controls aging and longevity of *C. elegans*, flies, and mammals (Kenyon, 2010). *C. elegans* is a powerful system to study the genetics of neuronal aging and regeneration (Ghosh-Roy and Chisholm, 2010; Hammarlund and Jin, 2014; Kenyon, 2010). Normally, when the DAF-2/IGF-1 receptor is activated it induces a conserved PI3K/AKT kinase cascade, which in turn inhibits the DAF-16/FOXO transcription factor from entering into the nucleus. Reduction of *daf-2* function doubles life span, and long-lived worms are considered to stay healthy for longer (Kenyon et al., 1993; Kenyon, 2010). In these animals, DAF-16/FOXO affects transcription of different genes that encode heat-shock proteins, antimicrobial proteins, antioxidants, and other molecules, which leads ultimately to extended lifespan (Lin et al., 2001; Murphy et al., 2003). Recently, aging-associated axonal morphological alterations and decline in regenerative capacity were described in *C. elegans*. *daf-2* mutations delayed these age-related morphological changes and improved regeneration of aged severed axons in a *daf-16-*dependent manner (Byrne et al., 2014; Pan et al., 2011; Tank et al., 2011; Toth et al., 2012). Moreover, in the case of GABA motor neurons, as adult animals age there is a reduction in axon growth, retraction and regrowth in response to injury. Surprisingly, the decline in regeneration by regrowth is controlled by *daf-16* in a cell-autonomously fashion and independently of lifespan (Byrne et al., 2014).

The process of regeneration of injured axons in vertebrates and invertebrates often involves degeneration of the distal part followed by regrowth (Chisholm et al., 2016; Park et al., 2008; Ruschel et al., 2015; Taylor et al., 2005; Yaniv et al., 2012). However, different invertebrates, including nematodes and crustaceans, use plasma membrane fusion as an alternative mechanism for repair of injured axons (Ghosh-Roy et al., 2010; Hoy et al., 1967; Neumann et al., 2015; Neumann et al., 2011). Cell fusion events were also observed in the brains of mammals both spontaneously and as a result of injury such as stroke (Alvarez-Dolado et al., 2003; Johansson et al., 2008; Paltsyn et al., 2013), but the role of these events has remained unclear (Giordano-Santini et al., 2016). In *C. elegans*, axonal regeneration via auto-fusion is mediated by the fusogen EFF-1 (Ghosh-Roy et al., 2010; Neumann et al., 2015; Neumann et al., 2011). EFF-1 is the first *bona fide* eukaryotic developmental cell-cell fusion protein. It is expressed in different cell types including neurons, and mediates fusion between cells by a homotypic mechanism (Gattegno et al., 2007; Mohler et al., 2002; Podbilewicz et al., 2006; Shemer et al., 2004). Thus, EFF-1 has to be expressed in both cellular membranes for them to merge and the high-resolution crystal structure of EFF-1 ectodomain suggests that EFF-1 acts by a mechanism that is distinct from viral-induced fusion. EFF-1, not only has to be present in both membranes for them to fuse, it also appears to interact in trans forming complexes that tether and fuse the membranes (Pérez-Vargas et al., 2014; Podbilewicz et al., 2006). During embryonic development EFF-1 is kept inactive inside early endosomes via Dynamin-and RAB-5-mediated endocytosis (Smurova and Podbilewicz, 2016).

We have been studying the role of fusion proteins in the PVD neuron, a polymodal nociceptor that senses harsh touch to the body, cold temperature, and posture (Chatzigeorgiou et al., 2010; Oren-Suissa et al., 2010; Smith et al., 2010; Tsalik et al., 2003). The PVD neuron exhibits an elaborate and invariant dendritic structure, which is composed of a repetitive unit that resembles a candelabrum (or a “menorah”) (Fig 1A) (Oren-Suissa et al., 2010). These dendritic trees arise and are dynamically maintained through several intrinsic and extrinsic genetic pathways (Chatzigeorgiou et al., 2010; Cohen et al., 2014; Dong et al., 2013; Dong et al., 2015; Husson et al., 2012; Liu and Shen, 2011; Maniar et al., 2012; Oren-Suissa et al., 2010; Salzberg et al., 2013; Smith et al., 2010; Wei et al., 2015).

**Fig 1.**
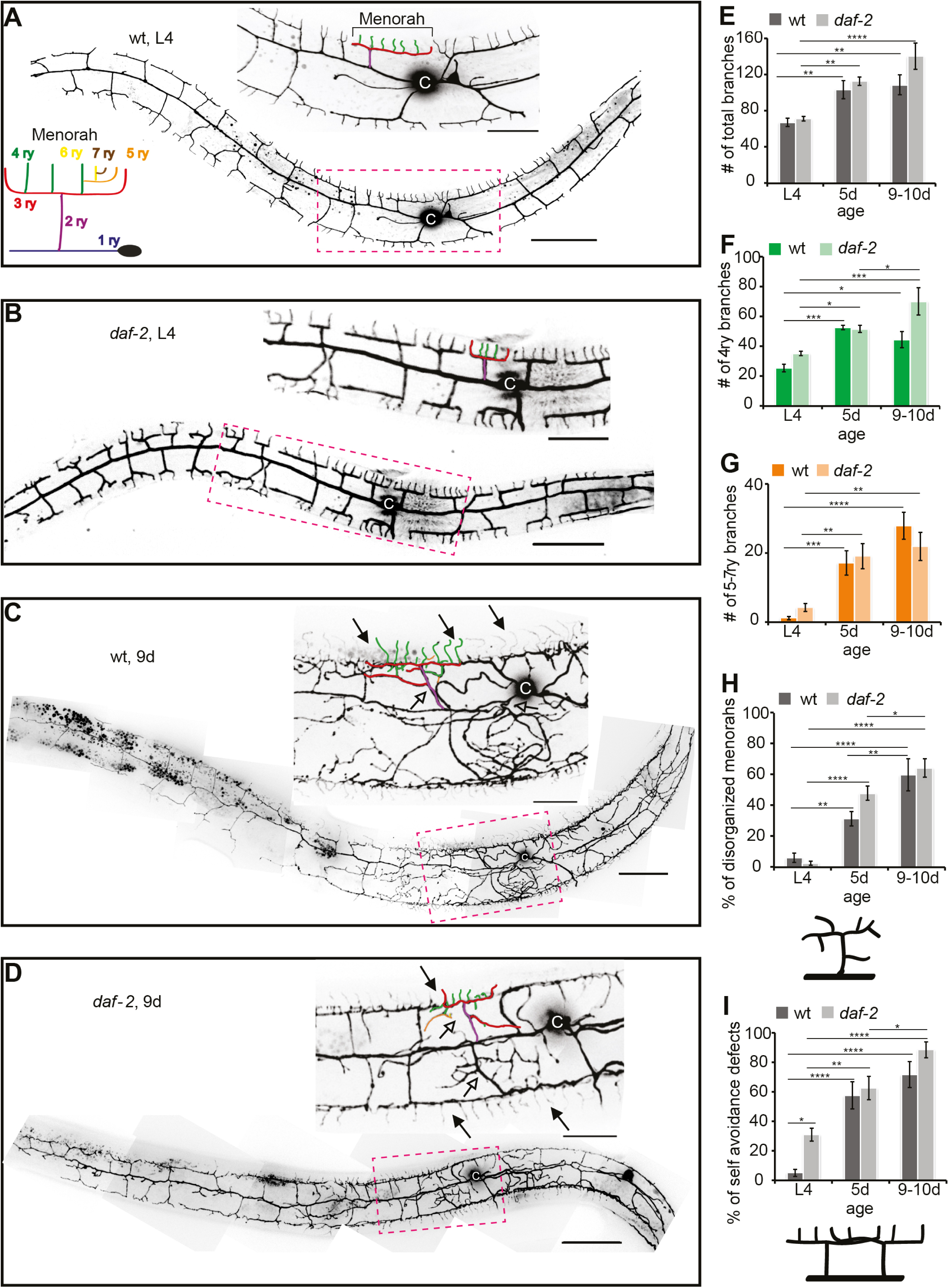
PVD’s dendritic alterations during aging. **(A-D)** PVD in wild-type (wt; **A** and **C**) and *daf-2* mutants (**B** and **D**) at L4 stage and 9 days of adulthood (9d). Upper panels, magnified boxed areas with candelabra-shaped units colored according to the scheme (**A).** c, cell body. Scale bars, 50 μm and 20 μm in magnified images. Anterior is left and ventral is down in all the figures. Filled arrows, self-avoidance defects. Empty arrows, disorganized menorahs. **(E-I)** Quantitation of phenotypes. Percentages were calculated for 100 μm of length around the cell body. Error bars, ± s.e.m. (**E**) There is no significant difference between wild-type animals and *daf-2* mutant animals at any age (one-way ANOVA). However, there is a significant difference between different developmental stages of the same genotype.(**F-G**) According to ANOVA age gives a significant difference in 4ry and in 5-7ry branching number both in wt and in *daf-2* backgrounds. (**H-I**) According to one-way ANOVA age has significance in menorahs disorganization and loss of self-avoidance in both wild-type and *daf-2.* Statistics were calculated using one way ANOVA followed by post-hoc analysis of student’s t-test with Benferroni correction. * *P<0.05, ** P<0.01, *** P<0.001, **** P<0.0001.* Number of animals analyzed: wt L4, *daf-2* L4 and *daf-2* 5d n=10, wt 5d n=9, wt 9d n=6, *daf-2* 9d n=8.

The PVD develops in a stereotypic fashion from the L2 larval stage to the young adult (Oren-Suissa et al., 2010). The fusion protein EFF-1 mediates dendrite retraction and auto-fusion of excess branches by a novel cell-autonomous pruning mechanism (Oren-Suissa et al., 2010). Thus, this fusogen maintains the structural units by trimming excess branching during normal developmental arborization (Oren-Suissa et al., 2010). AFF-1, a paralog of EFF-1, is a second *C. elegans* fusogen displaying a more restricted tissue distribution pattern (Avinoam and Podbilewicz, 2011; Sapir et al., 2007); AFF-1 was not found to be involved in PVD remodeling during development (Oren-Suissa et al., 2017). AFF-1 and EFF-1 fusion proteins also auto-fuse epithelial and myoepithelial cells to form tubes and reshape glial cells (Procko et al., 2011; Rasmussen et al., 2008; Soulavie and Sundaram, 2016; Stone et al., 2009). Moreover, it has recently been demonstrated that in vertebrates auto-fusion takes place in the development of the vascular endothelium, where it leads to pruning of excess blood vessels (Lenard et al., 2015) in a process that remarkably resembles EFF-1-mediated PVD pruning (Oren-Suissa et al., 2010) and that may link EFF-1 to the actin cytoskeleton. Recently it was reported that during larval development, EFF-1 interacts with filamentous actin via Spectraplakin (VAB-10A) to maintain EFF-1 at the site of epidermal cell fusion (Yang et al., 2017). The pathway of PVD dendritic repair following laser-induced dendrotomy and the functions of EFF-1 and AFF-1 were recently established (Oren-Suissa et al., 2017). The mechanism of PVD dendritic regeneration following injury can be divided in five overlapping stages: (1) reattachment at site of injury, (2) loss of self-avoidance between adjacent menorahs, (3) AFF-1-dependent menorah-menorah fusion that can bypass the lesions, (4) sprouting of compensatory dendrites and (5) EFF-1-dependent trimming of excess branches. It is unknown whether PVD structure and function is affected by aging and by the activity of fusion proteins.

Here, we show that induction of cellular fusion proteins (fusogens) can remodel and facilitate regeneration of dendrites in polymodal PVD neurons of aging *Caenorhabditis elegans*. Using whole-animal live imaging, we find that the PVD dendritic trees, composed of repetitive orderly multibranched units, show age-dependent hyperbranching, disorganization, and loss of self-avoidance. These processes, while independent of canonical lifespan-regulating pathways, can be partially rescued by ectopic expression of the fusogen EFF-1 that prunes multibranched dendrites. Surprisingly, most wild-type and *daf-2* (Insulin/IGF-1Receptor) mutant adults that have disorganized dendritic trees have normal response to harsh touch. Furthermore, we found a decreased capacity of old animals to repair laser-induced injured dendrites via auto-fusion that can be restored by reducing DAF-2 function or by ectopic expression of the EFF-1 paralog AFF-1 that remodels dendritic trees via merging of injured branches. Our findings demonstrate that fusogens are sufficient to maintain the dendritic arbor structure and increase its regeneration potential by auto-fusion in aging animals. These anti-aging strategies can be potentially applied to other organisms to help them recover from stroke, brain trauma and spinal cord injury.

## Results

### Progressive dendritic remodeling during aging is independent from the Insulin/IGF-1 pathway

Aging neurons lose their ability to remodel following injury, change their complex morphology, and may degenerate (Yankner et al., 2008). Whereas axon degeneration during aging has been studied extensively both in mammals and in invertebrates, the fate of complex dendritic trees in old animals has not been studied in detail in any organism. To determine how aging affects the complex arborized dendritic structure, we analyzed the *C. elegans* PVD dendritic branching patterns from the fourth larval stage (L4) to ten-day-old adults. We found that during aging PVD’s dendritic structures undergo disorganization and hyperbranching **(Figs 1A, 1C, and S1)**. Remarkably, we found that these age-dependent morphological changes of the PVD dendritic pattern were not affected in long-lived animals carrying a mutation in *daf-2* **(Figs 1A-1H, and S1)**. Adjacent PVD dendrites normally avoid each other (Smith et al., 2012), and we found that as the animal ages the structural units lose their self-avoidance properties (Fig 1C, 1D, **and** 1I). Consistently, this age-dependent dendritic pattern did not improve in *daf-2* mutant animals; young *daf-2* animals at the L4 stage exhibited significantly more *daf-16*-dependent self-avoidance deficiencies in comparison to wild-type (Fig 2). Thus, taken together our results reveal that during aging there is a *daf-2*-independent increase in disorganization of branching and loss of self-avoidance.

**Fig 2.**
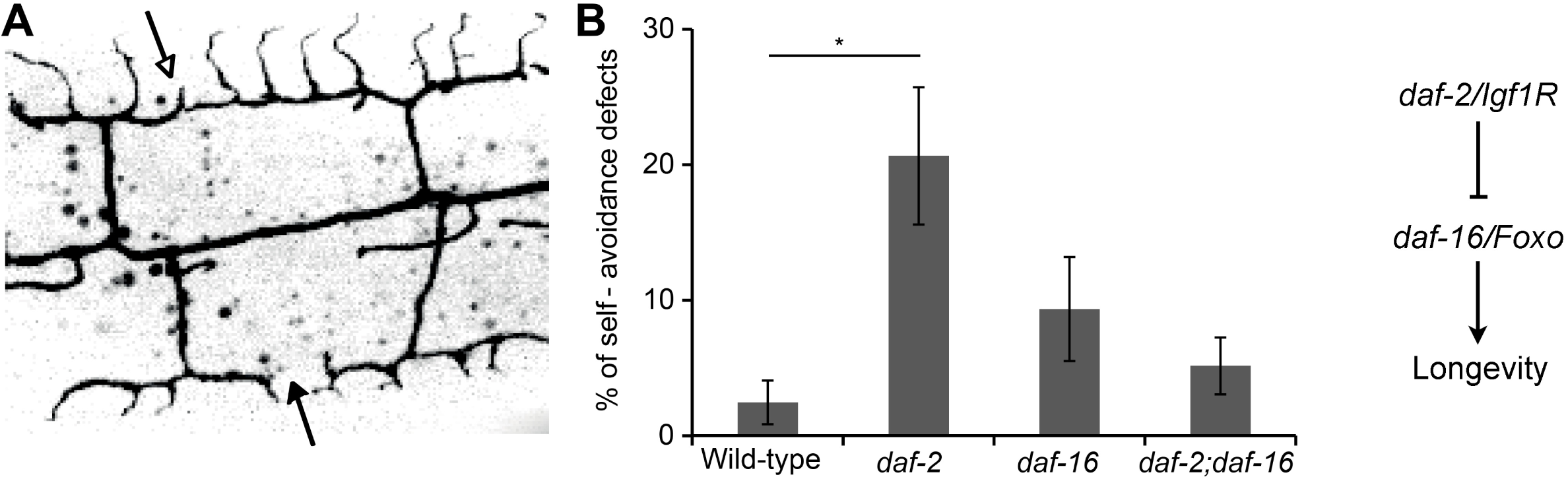
DAF-2-DAF-16 control self-avoidance at L4 stage. (**A**) Image showing four wild-type PVD dendritic units labeled with GFP, two of them do not overlap (filled arrow) and two show defective self-avoidance (empty arrow). (**B**) Percentage of defects in self-avoidance in 100 μm of length around PVD cell body at L4 stage in different genotypes, n ≥ 7. Defective self-avoidance increases in *daf-2* mutants in a *daf-16*-dependent manner. Error bars are ± s.e.m. Statistics were calculated using one way ANOVA followed by post-hoc analysis of student’s t-test with Benferroni correction. * P<0.05. Number of animals analyzed: wt n=7, *daf-2* n=12, *daf-16* n=11, *daf-2;daf-16* n=7

The elaborate PVD structure is highly dynamic in young animals, exhibiting constant growth and retraction of individual high order branches (Oren-Suissa et al., 2010; Smith et al., 2010; Smith et al., 2012). To further understand the dynamics of the age-dependent dendritic tree remodeling, we followed individual animals over time and analyzed time-lapse movies. We found that 5-day adults still exhibited some plasticity of the dendritic tree, with dynamic growth and retraction events (Fig 3A and **Movie S1)**; however, both growth and retraction were two times slower (Fig 3B) compared to younger L4 animals, imaged under the same conditions. Next we quantified the number of movements of branches. Most branches showed dynamic movements ending either with elongation or with shortening, but a minority of branches only grew or only retracted throughout the movies. When young animals were imaged, not only the rates of growth and retraction were higher, but also the number of dynamic events involving both growth and retraction were higher (data not shown). Interestingly, both in L4 larvae and 5d adults there were more events that ended with outgrowth or elongation rather than retraction events. We also found that the decrease in dynamic events with age was proportional both for outgrowth and retraction (Fig 3B). Thus, we conclude that at all ages there are slightly more growth events than retractions, which leads to branch accumulation with time and thus causing hyperbranched phenotype of the PVD. These experiments reveal that the PVD dendritic arbors grow and age in a dynamic way and not in a step-wise manner as apparent from still images. This dynamic growth and retraction together with the increase in branching as animals age suggest that hyperbranching is part of the normal dynamic growth of the dendritic arbors.

**Fig 3.**
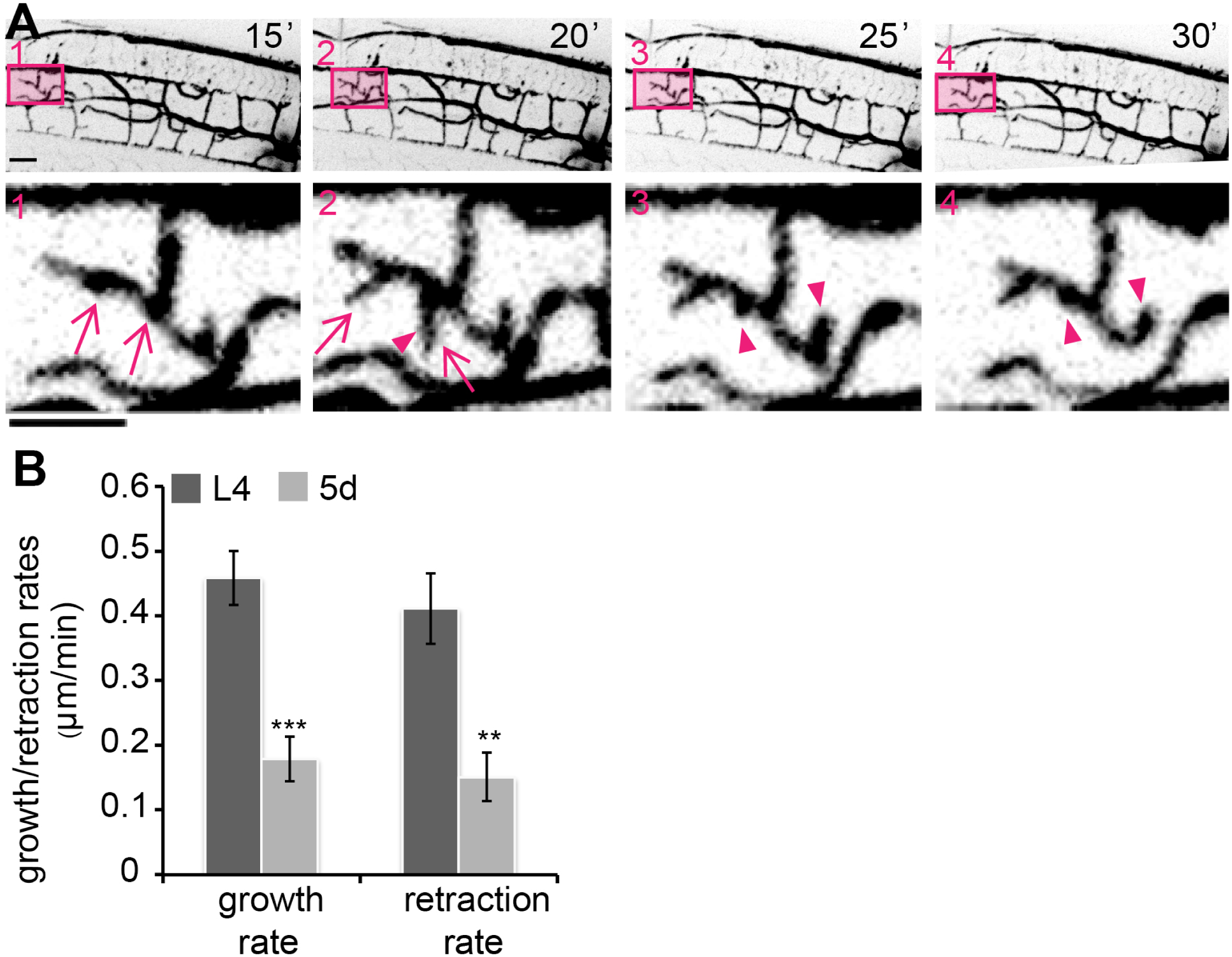
Adults show a reduction in dendritic plasticity. (**A**) Time-lapse confocal projections of 5 days of adulthood (5d) wild-type animal. Boxed areas are enlarged in the lower panel and reveal dynamic growth (arrows) and retraction (arrowheads) of branches. Scale bars, 10 μm. (**B**) Growth and retraction rates in μm/minute of branches at L4 stage and 5d as measured from time-lapse movies. ≥ 18 branches were analyzed from 4 L4 animals and 2 5d animals. Error bars are ± s.e.m. *P* values from *t* tests: ** *P<0.01,* *** *P<0.001*. See also **Movie S1**.

We speculated that reduced dendritic plasticity and structural changes in aged animals would affect the functionality of the PVD neurons. To shed more light on the link between structure and function of dendrites during aging, we performed a functionality test. PVD responds to harsh stimulations, and we used a classic harsh touch assay in *mec-4* mutant animals to specifically test PVD activity without the background light touch response mediated by the six light-touch mechanosensory neurons (Oren-Suissa et al., 2010; Way and Chalfie, 1989). First, we found that although *mec-4* animals have shorter life span (Pan et al., 2011), their dendrites looked similar to wild-type between the ages L4-5d (Fig 4), which further demonstrates that the morphological alterations we see are lifespan-independent. Second, we found that the PVD functionality showed a minor decrease with aging, with 5d adults presenting a small but significantly reduced harsh touch response in comparison to 1d adults, in both *mec-4* and *mec-4;daf-2* double mutants (Fig 4F). Harsh touch to the body was measured by prodding animals with a platinum wire in the midsection of the body (Oren-Suissa et al., 2010; Way and Chalfie, 1989). We cannot rule out that multiple components of the sensorimotor circuit contribute to the age-dependent decline in PVD activities. Thus, our results reveal that the morphological and behavioral hallmarks of aging in PVD dendritic arbors are independent from the canonical IGF-1 pathway that affects lifespan. Moreover, the fact that the dramatic increase in branching and loss of self-avoidance in middle-aged animals (5d-adults; Figs 1-3) correlates with lack of response to harsh touch in only 10% of the worms, compared to 1d adults, suggests that hyperbranching, loss of self avoidance and disorganization of dendritic trees might be normal and not deleterious for the behavioral response.

**Fig 4.**
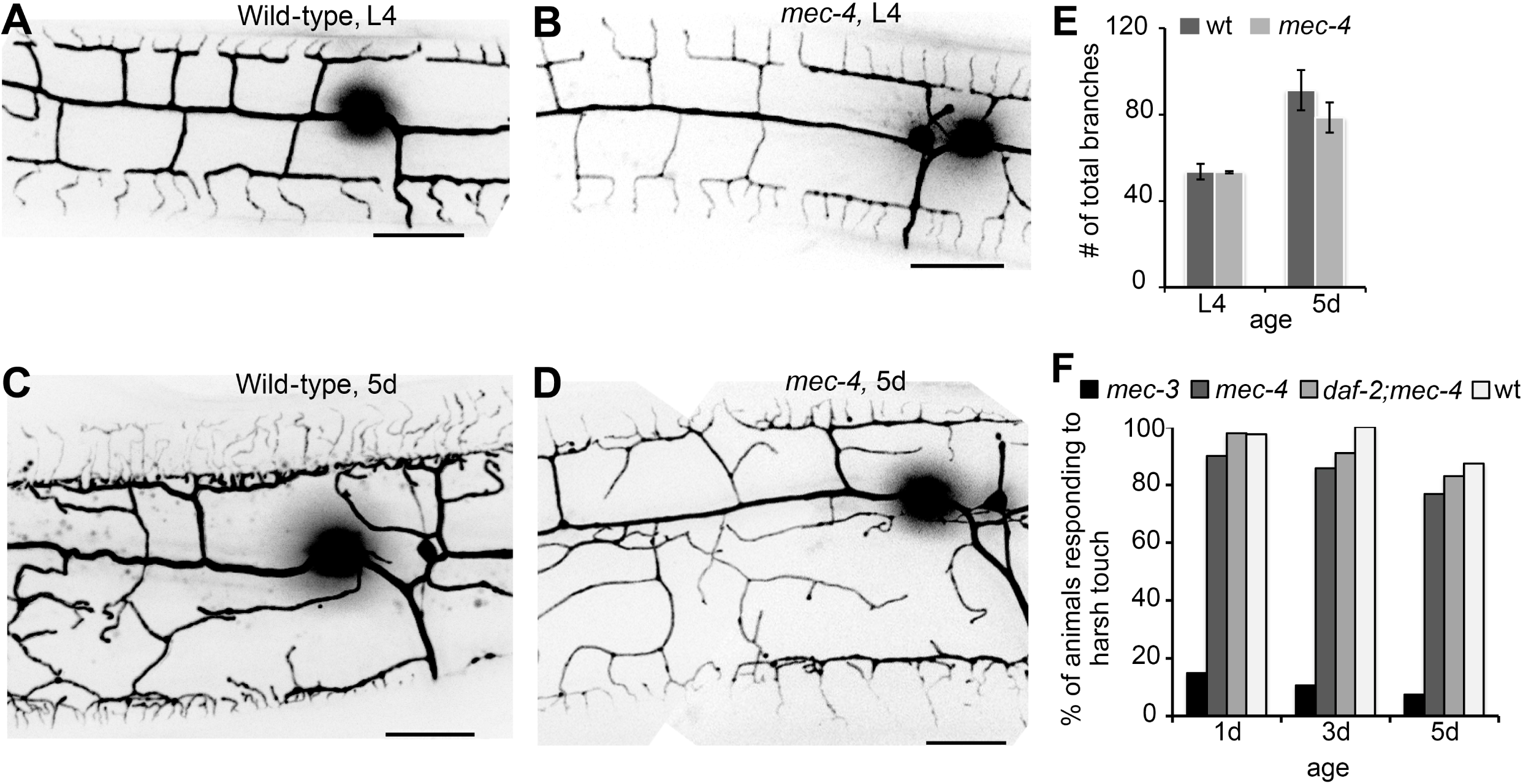
Decline in response to harsh touch during aging in both wild-type and *daf-2*. **(A-D)** PVD’s cell-body region of wt and light-touch insensitive *mec-4* mutants at the L4 stage and at 5 days of adulthood (5d). Scale bars are 20 μm. (**E**) Total number of branches counted in 100 μm of length around cell-body in wt and *mec-4* mutants at L4 and 5d (no significant differences between wt and *mec-4)*. Error bars are ± s.e.m. Number of animals analyzed: wt L4 and *mec-4* 5d n=7, wt 5d and *mec-4* L4 n=5 (**F**) Percentage of animals responding to harsh touch by escaping away from the stimulus. At the age of 5d, the response declined in *mec-4* light-touch defective mutants and in *daf-2;mec-4* double mutants. *mec-3* mutants are defective in harsh-touch mechanosensation and do not respond to harsh touch. We performed a one way ANOVA statistical test and found no significant difference between wt and *mec-4* animals (at any age), or between any of the backgrounds (chi square test). Number of animals analyzed: 1d *mec-3* n=88, *mec-4* n=70, *daf-2;mec-4* n=45, wt n=39; d3 *mec-3* n=57, *mec-4* n=70, *daf-2;mec-4* n=55, wt n=26; d5 *mec-3* n=68, *mec-4* n=134, *daf-2;mec-4* n=41, wt n=95.

In summary, the changes in dendritic architecture are not necessarily a deleterious aspect of aging, but could actually be part of normal neuronal maintenance that keeps the cell functioning over time. The fact that they are not altered in *daf-2* mutants supports this idea, as does the functional data that shows most of the neurons function fine in 5 days adults (Fig 4). Thus, the observed increase in dendrite branching and the disorganized appearance of the trees is probably adaptive as the animals increase in girth.

### Age-dependent dendrites remodeling can be modulated by EFF-1

The fusogen EFF-1 is essential to fuse some injured axons in *C. elegans* (Ghosh-Roy et al., 2010; Neumann et al., 2015), and is involved in PVD’s dendrite pruning in a cell-autonomous and dosage-dependent manner during development and in young adults (Oren-Suissa et al., 2010). When EFF-1 is overexpressed in the PVD, a strong gradient of arborization is seen in L4s and young adults, with almost complete lack of branches in areas that are distal from the cell body. However, in areas around the cell body the PVD menorahs appear similar to those in the wild-type (Oren-Suissa et al., 2010) (Fig 5A and 5B). Since EFF-1 retracts and simplifies dendritic arbors in young animals (Oren-Suissa et al., 2010) and is expressed in the PVD throughout adulthood (**Fig S2**), we hypothesized that EFF-1 overexpression will be able to retract dendrites in aged adults. Indeed we found that EFF-1 overexpression in the PVDs simplified the hyperbranching around the cell body at 9-10 days adults (Fig 5C-5E); in particular, the quaternary branch order was decreased (Fig 5F). A trend of reduction in fifth and higher order branches also appeared in aged animals overexpressing EFF-1 (Fig 5G). Thus, overexpression of the fusogen EFF-1 in the PVD neuron is sufficient to simplify aged menorahs. These results suggest that there is a reduction in EFF-1 expression in old animals, or a decreased efficiency of EFF-1 activity in older animals. We quantified the expression levels of EFF-1 in the PVD throughout development and found a reduction in the number of EFF-1 puncta in aged animals **(Fig. S3)**. In contrast to *eff-1*, *aff-1* mutants have no evident morphological phenotypes in the PVD and its expression has not been detected in this neuron (**Fig S2;** (Oren-Suissa et al., 2017)); moreover, when AFF-1 is ectopically expressed in the PVD in an *eff-1* mutant background it does not retract excess branching, demonstrating that it is unable to rescue the pruning defects of *eff-1* **(Fig S4)**. This result shows that EFF-1 and AFF-1 fusion proteins have different activities and are not interchangeable in the PVD neuron. In summary, during normal aging there is a reduction of EFF-1 expression that appears to result in an increase in dendritic branching. We can reverse the hyperbranching age-related phenotype by ectopic expression of EFF-1 in older animals, which simplifies the morphology of PVD dendritic arbors.

**Fig 5.**
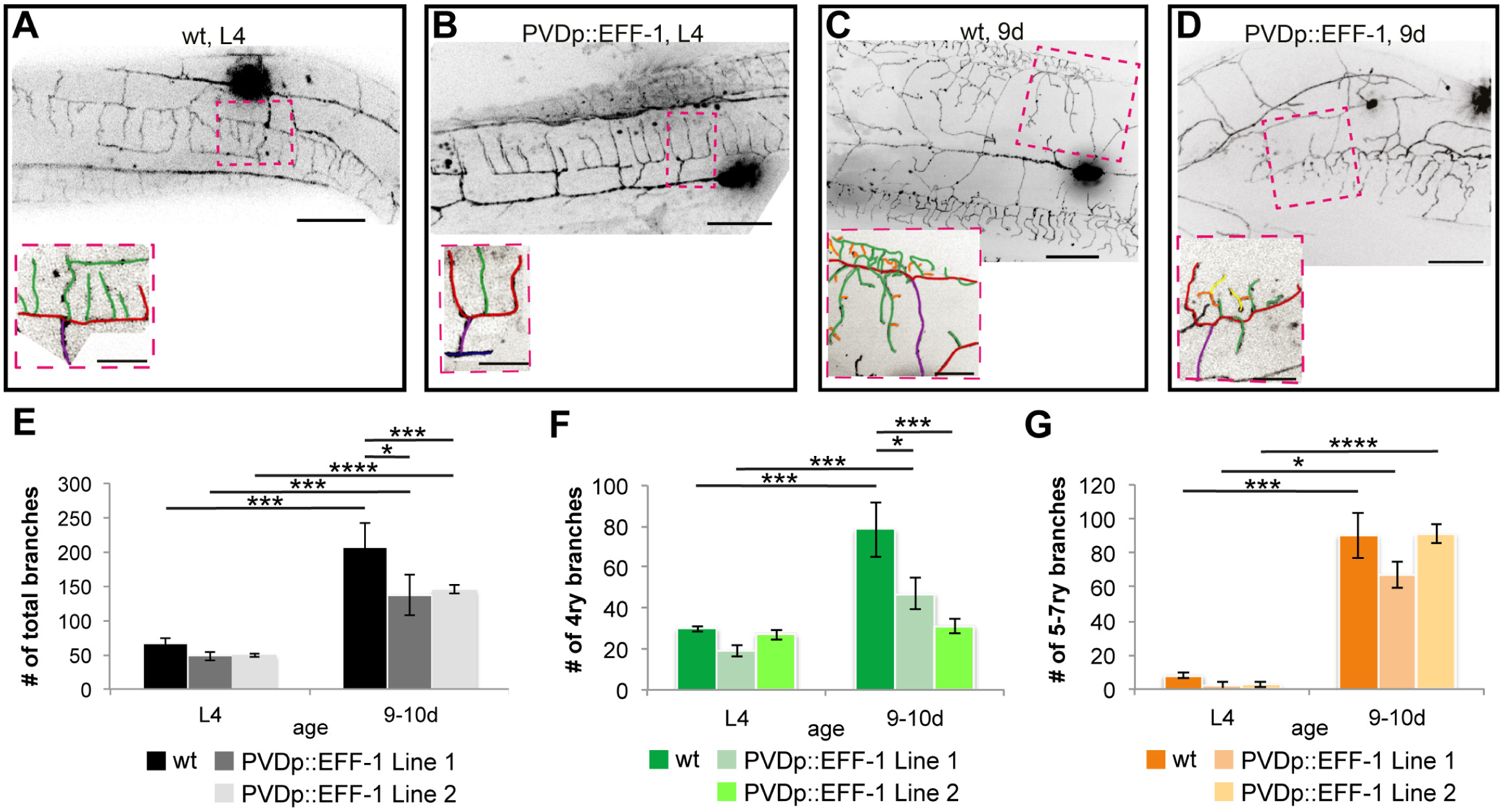
EFF-1 overexpression in the PVD simplifies aged dendritic trees. (**A-D**) Inverted fluorescence images of PVD neurons. (**A** and **C)** Represent wild-type neurons from L4 and 9 days of adulthood (9d). (**B** and **D)** Represent EFF-1 overexpression under PVD specific promoter (PVDp) at L4 and 9d. In each panel one candelabrum unit (boxed) is enlarged and colored (see Figure 1A). Scale bars, 20 μm and 10 μm in the enlarged images. **(E-G)** Graphs showing number of branches in 100 μm of length around cell body. Error bars, ± s.e.m. According to one way ANOVA test, age is a significant factor when comparing number of branches (*p<0.0001*). *p* values from *t* tests: * *p<0.05, *** p<0.001, **** p<0.0001* Number of animals analyzed: wt L4 n=8, wt 9-10d n=5, PVDp::EFF-1 line 1 L4 n=12 and 9-10d n=4, PVDp::EFF-1 line 2 L4 n=15 and 9-10d n=21. PVDp::EFF-1 line 1 and 2 correspond to worms carrying the extrachromosomal arrays *hyEx392* and *hyEx23,* respectively.

### Age-dependent decline in dendrite regeneration is dependent on Insulin/IGF-1

Mammalian axons regenerate better in younger individuals than in adults (Verdu et al., 2000). Similarly, axonal regenerative ability declines drastically as nematodes advance through development and age (Byrne et al., 2014; Zou et al., 2013). Our knowledge is still poor on the regenerative capacity of the dendrite, and how aging affects this process of neuronal repair (Bekkers and Hausser, 2007; Nawabi et al., 2012; Rao et al., 2016). To study dendritic regeneration we severed the primary dendrites of PVD neuron in aging adults. Typically, the PVDs show robust regeneration at L4 larval stage, consisting in dendrite sprouting from the proximal fragment still attached to the cell body and reconnection via fusion with the separated distal dendrite fragment (Fig 6A, **Movie S2**; (Oren-Suissa et al., 2017)). To directly measure reconnection by auto-fusion, we used the photoconvertible reporter Kaede **(Fig S5 and Movie S3)**. If fusion fails to occur, the detached distal part eventually degenerates. We found that 2-3 day-old wild-type adults respond more slowly to laser dendrotomy in comparison to L4 and young adults (∼70% of the young animals presented regeneration, whereas at the age of 2-3d neither regeneration nor degeneration occurred within 3-6 hours (Fig 6E **and Movie S4)**. At the age of 5 days, the ability to regenerate by dendrite auto-fusion was almost completely lost (Fig 6C and 6F). Thus, we found that relatively young adults already show a decline in their dendritic regenerative ability.

**Fig 6.**
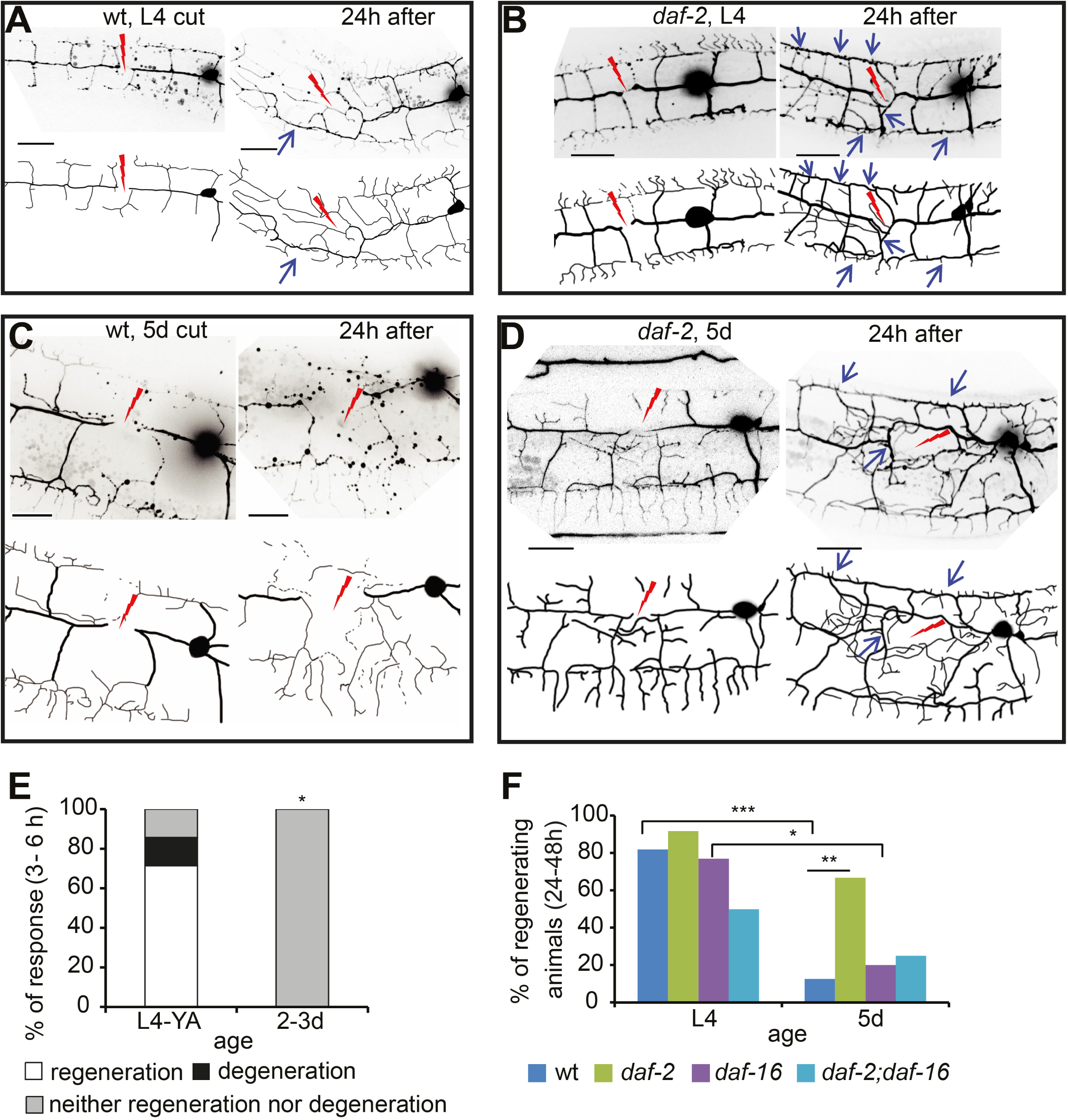
*daf-2*(-) restores regenerative ability of aged animals via *daf-16*. **(A-D)** PVD neurons immediately after cut and 24 h later in wild-type and *daf-2* mutants at L4 and 5 days of adulthood (5d), as indicated. Schematic illustrations below negative images. Red lightings, injury sites. Blue arrows, fusion sites (successful regeneration). Scale bars, 20 μm. (**E**) Response to injury of wild-types at L4-young adult (YA) and 2-3d within a short time (3-6 h) after injury. Number of animals: n=7 (L4 –YA) and n=4 (2-3d). (**F**) Percentage of successfully regenerating animals within 24-48 h post injury. Statistics were calculated using *Fisher’s exact tests* * *p<0.05*, ** *p<0.01, *** p<0.001.* Number of animals analyzed: wt L4 n=22, wt 5D n=16, *daf-2* L4 n=24, *daf-2* 5d n=12, *daf-16* L4 n=13, *daf-16* 5d n=10. See also **Movies S2-S4**.

To determine whether the decline in the ability to remodel dendrites by auto-fusion is dependent on the Insulin/IGF-1 pathway we tested whether the long-lived *daf-2* mutants are prevented from losing their dendritic self-fusion ability. We found that *daf-2* mutants showed similar regeneration to wild-type at L4 stage (Fig 6B and 6F), whereas at older age (5d) *daf-2* mutants had a much higher regenerative ability than wild-types (70% successful regeneration in *daf-2* versus 12.5% in wild-type) (Fig 6D and 6F). Similarly to axonal response to injury (Byrne et al., 2014), we found that DAF-2 inhibits regeneration of aged dendrites through inhibition of DAF-16, as *daf-16* mutants and *daf-2;daf-16* double mutants showed regenerative decline during aging similar to wild-type (Fig 6F). In conclusion, our results reveal that dendrite regeneration following transection declines with aging, a phenotype that is dependent on DAF-2/IGF-1R and its target DAF-16/FOXO.

### AFF-1 mediates and restores dendrite regeneration in aged animals

AFF-1 (Anchor cell Fusion Failure 1), is a transmembrane protein related to EFF-1 that executes several cell-cell fusion events during development in the epidermis, excretory, digestive and reproductive systems (Sapir et al., 2007). We have recently dissected the cellular pathway for dendritic remodeling following PVD injury (Oren-Suissa et al., 2017). We showed that injured primary dendrites regenerate and reattach at the site of injury via self-cell fusion. Simultaneously, terminal branches lose self-avoidance and grow towards each other, meeting and fusing via an AFF-1-mediated process. Throughout the regeneration process sprouting of compensatory branches occurs, and simplification of the dendritic arbors completes the regeneration process through *eff-1*-dependent pruning and retraction of excess branches. We found that AFF-1 functions cell non-autonomously from the epidermal seam cells and we hypothesize that AFF-1-containing extracellular vesicles fuse injured neuronal dendrites from without (Oren-Suissa et al., 2017).

Adults carrying loss-of-function mutations in *aff-1* have severe egg laying defects, shorter life spans and excretory system defects that prevented us from studying regeneration in aging *aff-1* mutant animals. To investigate whether AFF-1 can autonomously restore the regenerative ability of dendrites in older animals we overexpressed AFF-1 specifically in the PVD (PVDp::AFF-1; **Fig S6**). We found that the structural units of young animals expressing PVDp::AFF-1 appeared morphologically wild-type and responded similarly to dendrotomy **(Fig 7A, Movie S5)**. However, when 5-day old PVDp::AFF-1 animals were dendrotomized, the percentage of regenerating worms by dendrite fusion was significantly higher compared to wild-type animals (60-80% and 13%, respectively; Fig 7B **and** 7C, **Movie S6)**. Thus, AFF-1-specific overexpression in the PVD enables dendrite regeneration via auto-fusion of cut dendrites in older animals (Fig 7C).

**Fig 7.**
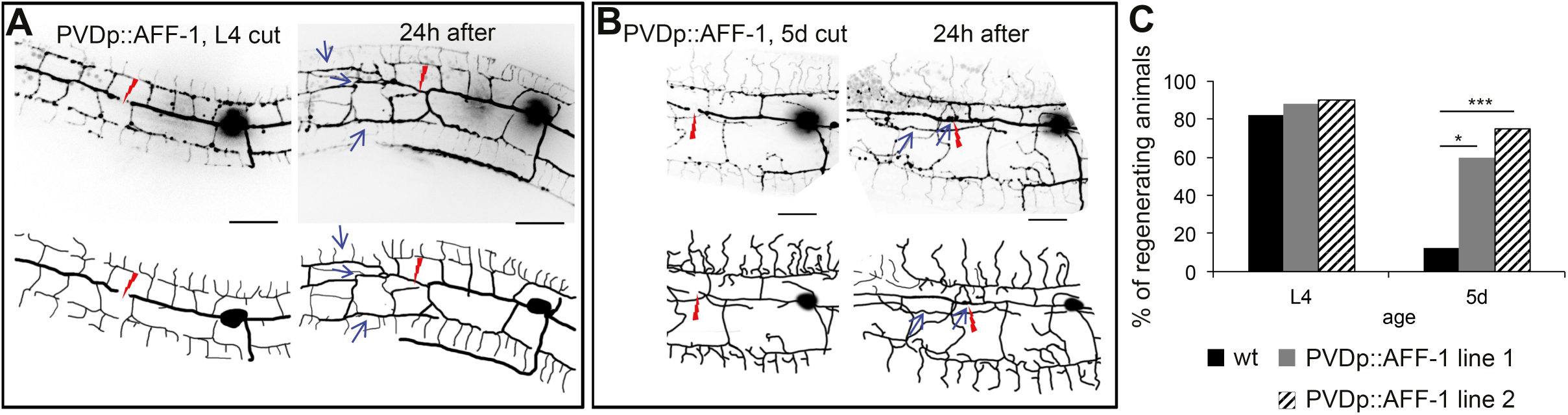
AFF-1 enhances regeneration by dendrite fusion in 5d-old animals. **(A-B)** Inverted fluorescence images of PVD neurons immediately after cut and 24 hours (h) later in animals overexpressing AFF-1 in the PVD (PVDp::AFF-1), at L4 and 5 days of adulthood (5d), as indicated on each image (schematic drawings below each image). Red lightning marks the injury site and blue arrows point at dendrite fusion sites. Scale bars, 20 μm. **(C)** Percentage of successfully regenerating animals within 24-48h after injury. *p* value from *Fisher’s exact test*: * *P<0.05; *** P<0.001*. Number of animals analyzed: wt L4 n=22, wt 5d n=16, PVDp::AFF-1 line 1 L4 n=17, 5d n=10, PVDp::AFF-1 line 2 L4 n=15, 5d n=11. See also **Movies S5-S6**. PVDp::AFF-1 line 1 and 2 correspond to worms carrying the extrachromosomal arrays *hyEx39* and *hyEx391,* respectively.

To determine whether *daf-2*(-) or PVDp::AFF-1 affect PVD architecture following injury independently of the remodeling by self-fusion, we quantified the number of PVD branches before and after injury in L4 and 5d adults. We found that only L4 animals expressing PVDp::AFF-1 showed an increase in the number of branches 24 hours post injury (Fig S7). Old 5d animals expressing PVDp::AFF-1 did not exhibit any difference in the number of branches as compared with wild-type and *daf-2* mutant animals, before and after injury. We also tested whether animals overexpressing AFF-1 have functional PVDs comparable to wild-type animals, by testing the behavioral response of PVDp::AFF-1 animals to harsh touch. We found that 80% PVDp::AFF-1 responding 5d adult animals (n=59), compared to control animals with 88% responding (n=75). These results suggest that there is no correlation between the ability to respond to harsh touch and the number of branches as a consequence of the expression of AFF-1 in the PVD. The main factor maintaining PVD activity is whether it regenerates via AFF-1-mediated auto-fusion following an injury, because it regenerates the complete dendritic tree structure. Taken together, the autonomous and ectopic expression of the fusion protein AFF-1 in the PVD appears to rejuvenate the ability to reconnect cut dendrites by the self-fusion mechanism. Since endogenous AFF-1 expression cannot be detected in the PVD ((Oren-Suissa et al., 2017); Fig. S2), it appears that ectopic activity of the AFF-1 fusion protein in the PVD neuron is sufficient to repair and remodel severed dendrites via the mechanism of auto-fusion in middle aged adults.

### Rejuvenation of fusion potential of cut dendrites by *daf-2(-)* and AFF-1(+)

We wished to further compare the regenerative modalities of AFF-1 overexpression and *daf-2* mutation following dendritic laser-induced severing in aging animals. Dendritic fusion following injury can occur by three possible pathways: 1) 3ry-3ry branch fusions that bypass the injury site (“menorah-menorah” fusion); 2) fusion of the proximal primary (1ry) branch to the detached distal 1ry; 3) both 3ry-3ry fusion and 1ry-1ry fusion (Fig 8A). To determine whether the mechanism of remodeling of dendritic trees following injury is distinct in wild-type, *daf-2* reduced function mutants and in animals ectopically expressing AFF-1 in the PVD, we compared the remodeling of dendritic trees in aging animals in these three genotypes following injury. We found that in wild-type and PVDp::AFF-1 animals at the L4 stage, the most prevalent mechanism of repair is 3ry-3ry fusion (pathway 1), whereas in *daf-2* mutants 3ry-3ry fusion together with 1ry-1ry fusion (pathway 3) increased compared to wild-type (Fig 8B-8D). In wild-type, we found degeneration in 50% (2-4d) and 80% (5-6d) of the dendrotomized animals (Fig 8E and 8H). Adult *daf-2* mutant animals (5-6d) presented a response to injury that resembled that observed in L4 wild-type animals, with 70% of the animals showing regeneration, mainly via 3ry-3ry fusions (Fig 8F and 8I). In contrast, in PVDp::AFF-1 animals the response to injury followed a different mechanism, with more regeneration via enhanced 1ry-1ry fusion instead of 3ry-3ry fusion as observed in *daf-2* mutants (Fig 8G and 8J). However, the AFF-1 effect was unrelated to longevity as PVDp::AFF-1 animals had normal lifespans **(Fig S8)**. Thus, differential activities of AFF-1(+) and *daf-2*(-) maintain the remodeling potential of aging arbors, and the mechanisms of rejuvenation by fusion that prevent degeneration appear to be independent. In both cases degeneration of the distal trees is prevented.

**Fig 8.**
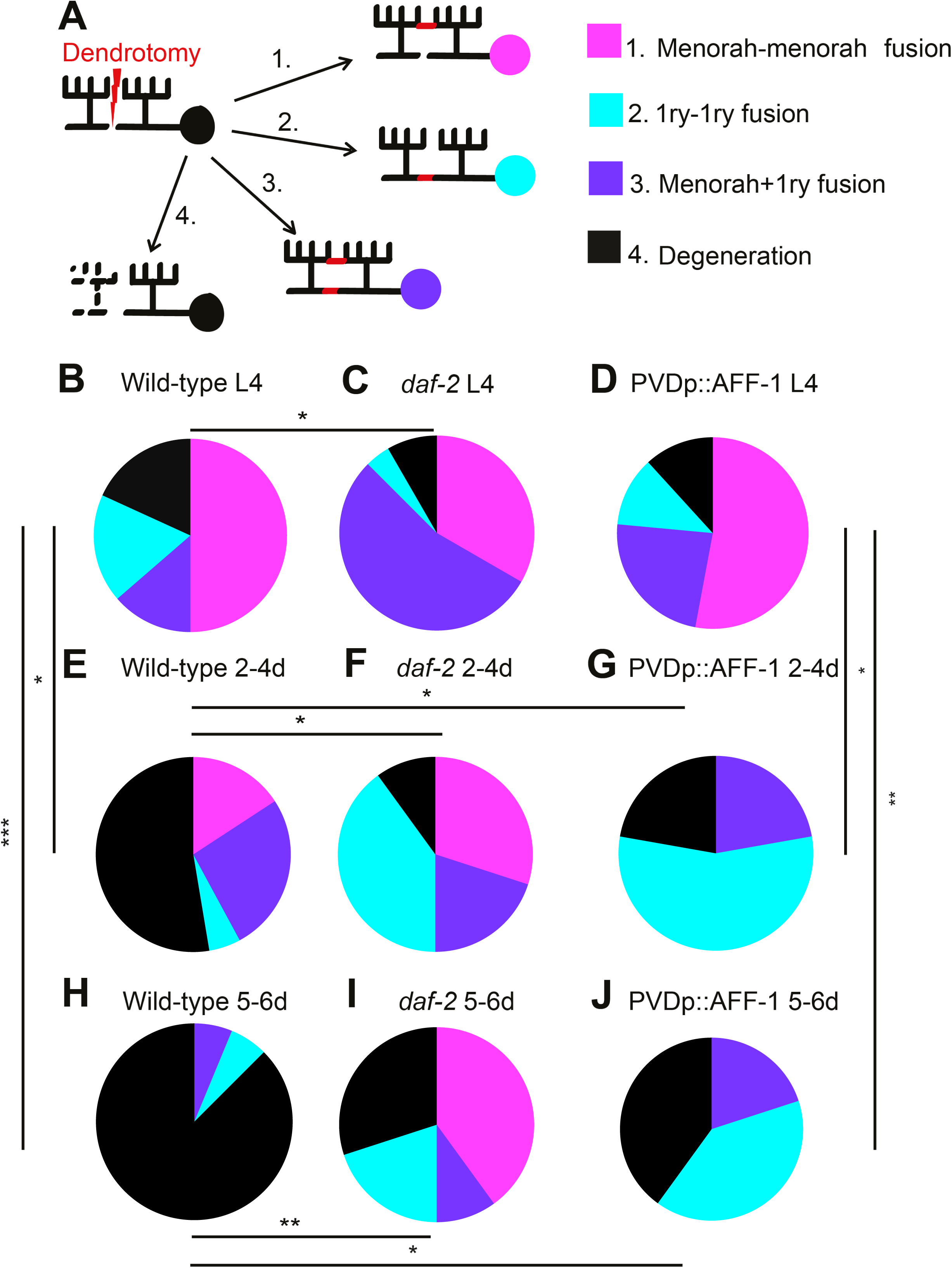
*daf-2*(-) and AFF-1(+) rejuvenate fusion potential of broken dendrites. (**A**) Cartoon representing four possible outcomes of dendrotomy. Dendrite auto-fusion indicates successful regeneration (outcomes 1-3), if fusion does not occur degeneration takes place (outcome 4). **(B-J)** Percentages of wild-type worms (wt), *daf-2* mutants and AFF-1 overexpressing animals (PVDp::AFF-1) at different ages, divided into the four different types of response to injury as described in (**A)**. Genotype and age are listed above each plot. *p* values from *Fisher’s exact tests*: * *p<0.05*, ** *p<0.01, *** p<0.001.* Additional significant differences were found between: (**C** and **F)**, (**C** and **I)** and between (**B** and **J)** (*p<0.05*).

Aged dendritic trees lose their regenerative potential because following laser-induced dendrotomy old neurons usually fail to auto-fuse broken dendrites and undergo degeneration. DAF-2/ IGF-1R negatively regulates the regeneration process. When the fusogen AFF-1 is ectopically expressed in the PVD neuron it has an antiaging activity that promotes auto-fusion of old transected primary dendrites.

## Discussion

Precise dendritic arborization is critical for proper functioning of neuronal networks, while defective dendritic development and maintenance lead to neuropathologies (Jan and Jan, 2010; Koleske, 2013). There is evidence from mammalian CNS showing increase in number and length of terminal dendritic arbors during aging, and reduced arborization in senile dementia (Buell and Coleman, 1979). However, it remains unclear whether there is a direct link between altered dendritic structure and reduced function in mammals and other organisms (Maletic-Savatic et al., 1999; Thompson-Peer et al., 2016). Axons of aging neurons also show altered morphologies in diverse species. In *C. elegans* these axonal alterations are not affected by organismal longevity (Byrne et al., 2014; Hammarlund et al., 2009; Verdu et al., 2000; Zou et al., 2013). Here we also found that dendritic alterations during aging are not affected by chronological age; however, unlike the PVD dendrites, in axons the *daf-2* mutation was found to delay the morphological alterations of aged animals (Pan et al., 2011; Tank et al., 2011; Toth et al., 2012). Nevertheless, the *daf-2* effect in age-related axonal branching was reported to be uncoupled of its role in extending the lifespan of the worms (Tank et al., 2011; Toth et al., 2012). It thus appears that DAF-2 is specifically involved in cell-autonomous pathways that maintain axonal, but not dendritic morphology. We observed a small but significant decline in response to harsh touch during aging in both wild-type and *daf-2* (Fig. 4). Thus, we propose that the dendrite overgrowth observed in old animals reveal a reduction in morphological plasticity and a normal change in the architecture of the dendritic trees that do not appear to have major functional consequences in the ability to respond to noxious mechanosensory stimuli. However, the link between structure and function needs to be further investigated in future studies. While progressive age-related hyperbranching of the PVD arbors was independent of insulin/IGF-1 pathway, overexpression of EFF-1 in the PVD was sufficient to partially rescue this phenotype in old animals (Figs 5 **and** 9). EFF-1 appears to act cell-autonomously to simplify the dendritic trees via its pruning activity mediated by branch retraction (Oren-Suissa et al., 2010).

**Fig 9.**
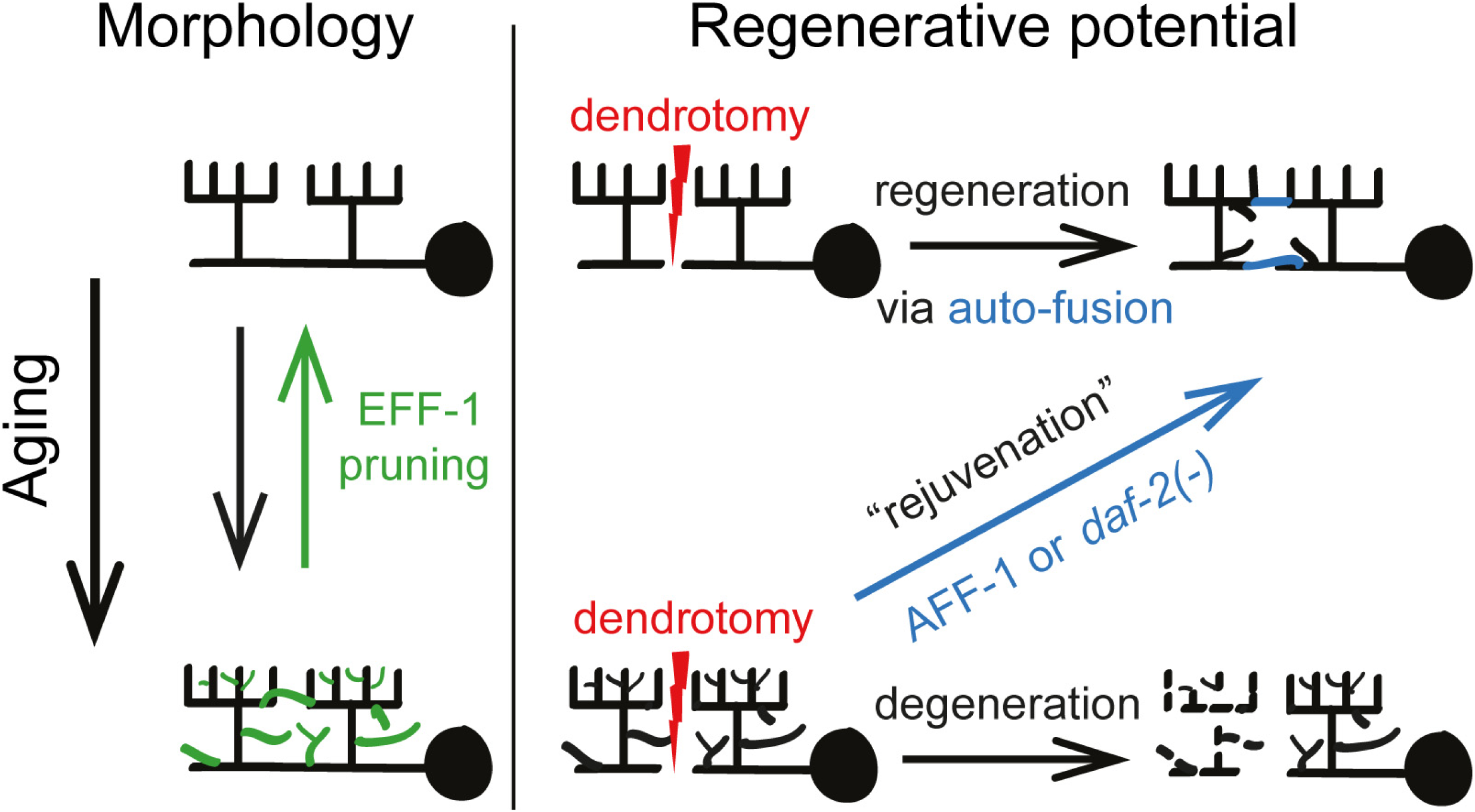
Model of morphological and regenerative antiaging activities on dendritic arbors. ***Morphological aging*** of PVD menorahs is a progressive and dynamic process that results in loss of self-avoidance, disorganization and hyperbranching. When the fusogen EFF-1 is ectopically expressed in the PVD neuron it has an antiaging activity that involves pruning of hyperbranched dendritic trees. **Aged menorahs lose their *regenerative potential*** because following laser-induced dendrotomy old neurons usually fail to auto-fuse broken dendrites and undergo degeneration. DAF-2/ IGF-1R negatively regulates the regeneration process. When the fusogen AFF-1 is ectopically expressed in the PVD neuron it has an antiaging activity that promotes auto-fusion of old transected primary dendrites.

We further found that dendrite regeneration following PVD dendrotomy decreases with age. This decline was “rescued” in *daf-2* mutants (Fig 6). Thus, repair of injured dendrites is *daf-2*-dependent whereas the hyperbranching is not (Figs 1, 6 **and** 9). Axonal regeneration is known to decline with age in different organisms (Byrne et al., 2014; Gabel et al., 2008; Hammarlund et al., 2009; Pestronk et al., 1980; Verdu et al., 2000; Zou et al., 2013). In GABA neurons of *C. elegans* the decline in axonal regeneration is also delayed in *daf-2* mutants (Byrne et al., 2014). However, these neurons do not engage in self-fusion during regeneration, whereas in PVD dendrites the main outcome of regeneration is auto-fusion. Recently, an important role of auto-fusion during axonal regeneration was demonstrated in the PLM sensory neurons of *C. elegans*, a process mediated by the fusogen EFF-1 (Ghosh-Roy et al., 2010; Neumann et al., 2015). In the PVD EFF-1 is not necessary for dendrite reconnection following injury (Oren-Suissa et al., 2017). In the PVD dendrites, this role is fulfilled by the EFF-1 paralog AFF-1. Significantly, when AFF-1 is ectopically expressed, this fusogen plays a role in maintenance of regenerative potential during aging (Figs 7-9). These findings highlight fusion mechanisms as potential target for pharmacological intervention in neuropathologies that result from both injury and aging.

## Materials and Methods

### Nematode strains

Animals were maintained at 20°C, unless otherwise stated, according to standard protocols (Brenner, 1974). All experiments were performed on hermaphrodites. N2 served as wild-type, and the following mutants and transgenic strains were used:

NC1841 *[wdIs52(F49H12.4::GFP); rwIs1(pmec-7::RFP)]* (Smith et al., 2010), outcrossed 4X, CHB392 *[hmnEx133(ser-2prom3::kaede)],* kindly provided by Candice Yip and Max Heiman (Yip and Heiman, 2016), BP709 *[hmnIs133 (ser-2prom3::kaede)]*, kindly provided by Tamar Gattegno, MF190 *[hmIs4(des-2::gfp, pRF4)]* (Oren-Suissa et al., 2010). CB1338 *[mec-3(e1318)IV]*, *Caenorhabditis* Genetics Center (CGC), CB1611 *[mec-4(e1611)X],* (CGC), BP176 *[hyEx23(des-2::eff-1, des-2::gfp, pRF4)]* (Oren-Suissa et al., 2010), BP177 *[hyEx392(des-2::eff-1, des-2::gfp, pRF4)],* BP488 *[eff-1(hy21)II; hmIs4; hyEx39(des-2p::AFF-1,myo-2::GFP,KS)]*, BP906 *[daf-2(e1370)III; wdIs52; rwIs1],* BP911 *[hyEx39; hmnEx133]*, BP915 *[hyEx39; hmnIs133]*, BP550 *[hyEx391(des-2p::AFF-1, myo-2::GFP, KS); hmnEx133],* BP919 *[daf-2(e1370)III; daf-16(mu86)I; hmnIs133],* BP923 *[daf-16(mu86)I; hmnIs133]*, BP924 *[daf-2(e1370)III; hmnIs133]*, BP925 *[mec-4(e1611)X; hmnIs133],* BP926 *[daf-2(e1370)III;mec-4(e1611)X; hmnIs133],* BP1056 *hyEx68[AFF-1::TY1::EGFP::3xFLAG, pRF4, KS]; dzIs53[F49H12.4p::mCherry], AFF-1::TY1::EGFP::3xFLAG* was kindly provided by transgenome project (Sarov et al., 2006), BP1055 *hyEx66[AFF-1::TY1::EGFP::3xFLAG, pRF4, KS]; dzIs53,* BP500 *zzIs22 [pJdC41(eff-1p::EFF-1::GFP), pRF4]; dzIs53 (pF49H12.4::mCherry) zzIs22* was kindly provided by William Mohler (del Campo et al., 2005).

### Molecular Biology

The *ser-2prom3::Kaede* (pCY2 plasmid) is a kind gift from Candice Yip and Max Heiman (Yip and Heiman, 2016). It was constructed by digesting *ser-2prom3* with SbfI and AscI and Kaede with AscI and NotI and then ligated together.

The *des-2p::AFF-1* construct (pME4 plasmid) was constructed by inserting an AFF-1 genomic fragment, digested from *hsp16.2::AFF-1* (Sapir et al., 2007) with NheI and KpnI and cloned into pME2 (Oren-Suissa et al., 2010) cut with the same enzymes. *des-2p::AFF-1* was injected at a concentration of 0.1ng/ml into *eff-1(hy21)* animals and three independent lines were obtained. For dendrotomy experiments, two lines (*hyEx39* and *hyEx391*) were crossed into a wild-type background expressing the PVD marker *ser-2p3::Kaede.*

### Live imaging of worms

Time-lapse imaging and short imaging (one time point) of worms by Nomarski optics and fluorescence microscopy was performed using a Nikon eclipse Ti inverted microscope with Yokogawa CSU-X1 spinning disk or using the Zeiss Laser Scanning Microscope (LSM) 510 META. Animals were anesthetized using 0.1% tricaine and 0.01% tetramisole in M9 solution for 20-30 minutes, and then they were transferred to a 3% agar slide with an eyelash attached to a toothpick. For short time imaging worms were often mounted on 3% agar slides containing 5–10 mM NaN_3_ instead. Image acquisition was done using Andor iQ or Metamorph software, when using the spinning disk confocal (SDC), and Zen software when using the LSM 510 meta microscope. Z-stacks were taken with PlanApochromat 60x oil NA=1.4 objective using the SDC or 63x NA=1.4 objective using the LSM. Excitation of GFP and green Kaede was done with 488 nm wavelength laser and 525 filter (6-15%, 50 ms exposure time), RFP and red Kaede was excited with 561 nm wavelength laser and 607 filter (15-20%, 50-100 ms exposure time). When using the sCMOS (Andor) camera z-stacks were taken with ∼0.23 μm z-step. With iXon EMCCD camera (Andor) z-stacks were taken with ∼0.5 μm z-step, gain was ∼100. When the LSM 510 meta was used, z-step was ∼0.8 μm. Multidimensional data was reconstructed as maximum intensity projections using the FIJI software (NIH Image). Images were prepared using Imaris, FIJI and Adobe Photoshop CS5. Final figures were constructed using Adobe Illustrator CS5.1.

### Laser dendrotomies

Micropoint, pulsed nitrogen laser, or the tunable Chameleon Ultra Ti-Sapphire laser system (two-photon), were used to sever the primary dendrite of the PVD. The Micropoint system was used on the Nikon eclipse Ti inverted microscope with Yokogawa CSU-X1 spinning disk. In order to transect neurons, we used 405 beamsplitter and 365nm dye cell and either IQ or Metamorph software. We used highest levels with the attenuator plate (all the way in), while the software controlled attenuator was adjusted between 80-90% when using IQ, and 30-50% when using Metamorph. Roughly 15 pulses at 10 Hz were administered for each cut in IQ and Metamorph. For all worms the primary dendrite was injured anterior to cell body. Animals were imaged and a z-stack was collected immediately after cut to confirm that the injury was successful.

The two-photon system was used on the Zeiss LSM 510 META microscope. The laser produces 200fs short pulses at 113MHz repetition rate and energy of 5nJ. In order to cut neurons we used 820 nm wavelength and 20-30% laser power using the Zen software. Worms were imaged immediately after cut to confirm that the injury was successful, and z-stacks were taken using the spinning disk confocal system or the LSM 510 META as described for live imaging of worms. After surgery, animals were recovered on NGM agar plates seeded with OP50 in a drop of M9 and imaged again later, or time-lapse movies were immediately acquired.

Regeneration was defined as continuation of the fluorescent signal between the distal and proximal ends or using Kaede photoconversion (**Fig S5**; see below). Significant differences between ages and genotypes were determined using *Fisher’s exact test.*

### Kaede photoconversion

In order to verify that dendrites fuse as response to injury we used the photoconvertible protein Kaede driven by a PVD specific promoter *ser-2prom3* (Yip and Heiman, 2016). Irreversible photoconversion of the green Kaede to red Kaede was achieved using Mosaic system on the Nikon eclipse Ti inverted microscope with Yokogawa CSU-X1 spinning disk, at 405 nm, 20-50 ms exposure time with 10-20 repeats across the region of interest, which was always the cell body, using either IQ or Metamorph software.

### Morphological quantitation of the PVD

Branch count and disorganization of PVD structural units (menorahs) was counted in the 100 μm around cell body (unless otherwise stated), as previously described (Oren-Suissa et al., 2010). Lack of self-avoidance between adjacent menorahs was determined in the same region as previously described (Smith et al., 2012). Z-stack maximum intensity projections of time-lapse movies were analyzed manually, using FIJI software. Factorial analysis of variance (ANOVA) and *t-tests* were performed to compare between genotypes and ages.

### Harsh touch assay

Harsh touch assay was performed as previously described (Oren-Suissa et al., 2010; Way and Chalfie, 1989) and the experimenter was blind to the genotype. The experiments were done in light touch mutant background (*mec-4(e1611)*) (Chalfie and Sulston, 1981). *mec-3(e1338)* worms were used as negative control (Way and Chalfie, 1989). These mutants are defective in harsh-touch mechanosensation and do not respond to harsh touch. Significant differences between the ages were determined by the *χ* 2 test.

### Life-span assay

Lifespan assays were carried out at 20°C as previously described (Apfeld and Kenyon, 1998). A population of worms was synchronized using hypochlorite and NaOH solution and ∼100 animals at the L4 “Christmas tree” stage were placed on NGM plates containing 49.5 μM 5-Fluoro-2'-deoxyuridine (FUDR, Sigma, F0503) seeded with OP50 *E. coli*, at a density of 20 worms per 6 cm plate. Adult day 1 was designated as the day after the L4 larval stage that served as time 0. Animals were scored as dead or alive every 2-3 days until all animals died. An animal was scored dead when it did not move or react at all to prodding with a platinum wire. Animals that crawled off of the plate, were trapped in fungal infection, “exploded” (i.e., showed extruded tissue from the vulva), or showed internal hatching (“bagging”), were excluded. If plates were contaminated, animals were transferred to fresh plates, using platinum wire. Statistical significance was determined using the Log-Rank (Mantel-Cox) test.

### RNA Isolation, Reverse Transcription, and Quantitative Real-Time PCR

Total RNA from Wild-type worms and des-2p::AFF-1 transgenic worms was isolated using the RNeasy Micro kit following the manufacturer’s protocol (Qiagen). RNA was used for cDNA synthesis using the qScript cDNA Synthesis Kit (Quanta BioSciences) with random primers. Real-time PCR was performed using AFF-1 and Fast SYBR® Green Master Mix (Applied Biosystems). The PCR amplification conditions were as follows: 40 cycles of 95 °C for 20 s and 60 °C for 20 s. Each PCR was run in triplicate. Data analysis and quantification were performed using StepOne software V2.2 supplied by Applied Biosystems. To account for the variability in the initial concentration of the RNA, results were normalized to the ACT-1 housekeeping gene.

### Statistical analysis

Statistical analysis was performed using the GraphPad online tool: http://www.graphpad.com/quickcalcs/ for *t-tests, Fisher’s exact test*s and *χ* ^2^ tests. Analysis of variance (ANOVA) that was performed when more than two groups were compared and Log-Rank survival test, using SPSS. *Freeman-Halton* extension of the *Fisher’s exact test* (for 2x4 contingency table) was performed using the Vassarstats.net online tool, to compare between the distributions of response to injury between different genotypes and ages.

## Acknowledgments

We thank D. Cassel, M. Hilliard, and A. Sapir for critically reading earlier versions of the manuscript. M. Heiman and C. Yip for providing CHB392, C. Smith and D. Miller for NC1841, and T. Gattegno for BP709. We also want to thank R. Kishony and all members of the Podbilewicz laboratory for discussion. Some strains were provided by the CGC, which is funded by NIH Office of Research Infrastructure Programs (P40 OD010440).

The authors declare no conflict of interest.

### Author contributions

VK, MOS and BP designed the experiments. VK performed most experiments. MOS developed the system to study neuronal degeneration and regeneration following dendrotomy using the PVD neurons. MOS generated the PVDp::AFF-1 and PVDp::EFF-1 transgenics and MOS and VK tested the effects of EFF-1 and AFF-1 over-expression on regenerating animals. BP did some harsh touch assays and supervised this work. VK, MOS and BP wrote the paper.

This article contains supporting information including eight figures and six movies

### Funding

This work was supported by the Israel Science Foundation (ISF grant 443/12, to B.P.), the European Research Council (Advanced grant ELEGANSFUSION 268843 to B.P.) and the HFSP long-term postdoctoral fellowship (LT000727/2013L to M.O.S.).

